# Reconstructing a physiological state space via chronic jugular microdialysis in freely moving mice

**DOI:** 10.64898/2025.12.08.692974

**Authors:** Michele Nardin, Nan Wang, Soad Elziny, Claire Boyer, Vojko Pjanovic, Luisa Schuster, Peter Boklund, Sarah Lindo, Kendra Morris, Anoj Ilanges, Jakob Voigts, Emily Jane Dennis

**Affiliations:** Janelia Research Campus, Howard Hughes Medical Institute, Ashburn, VA, USA; CMA Microdialysis AB, Kista, Sweden

## Abstract

Maintaining physiological homeostasis requires a complex interplay among endocrine organs, peripheral tissues, and distributed neuroendocrine control circuits, all of which are coupled through feedback loops that operate over minutes to hours. Although many physiological needs are broadcast through hormones, metabolites, and other chemical compounds circulating in the bloodstream, we rarely observe more than a few of these messengers together and at high cadence during behavior. To address this, we developed a minimally disruptive workflow to measure the free fraction of hundreds of amines and small peptides at a 7.5-minute cadence for ∼8 hrs in freely moving mice using chronic jugular microdialysis implants and chemical isotope labeling Liquid Chromatography-Mass Spectrometry. Single-compound profiles behave according to known physiology, such as purine turnover correlating with movement, delayed histamine/5-HIAA changes, and coordinated amino-acid dynamics. Our multiplexed measures enable high-dimensional analyses that uncover properties of the underlying dynamics. For example, systems-level analyses show that 10 dimensions explain over 70% of the variance in hormone/metabolite covariation, consistent with a low rank description of the physiological state space, with projections aligned to locomotion state transitions. Our work opens avenues for the discovery of hormonal dynamics, compound interactions, and their effects on behavior.

## Introduction

Physiological homeostasis is critical for animal survival. In mammals, bloodborne messengers act as a body-wide broadcast medium, traveling through the bloodstream to signal physiological needs to cells throughout the body (1–3). Homeostasis is regulated by control circuits in the hypothalamus, brainstem, and autonomic ganglia (4–13). For example, orexin cells in the lateral hypothalamus respond to the slope of glucose changes rather than its absolute values, and help mediate glucose-dependent decrease in locomotion, suggesting that the dynamics of blood chemistry are important signals to the brain and behavior (14). Despite the importance of these functions, the study of multiple bloodborne messenger dynamics during freely moving, in vivo animal studies is still challenging (Supp. Table 1).

Measuring many bloodborne molecules in vivo during free behavior presents a technical challenge (15). Unlike glucose monitoring, which is performed with simple electrochemical methods, many bloodborne molecules, such as amines, peptides, proteins, and steroid hormones, necessitate specialized measurement tools and often require blood sampling (16). Restraining animals for repeated collections can increase animal stress (17), and whole-blood draws are limited in volume and frequency to prevent hypovolaemia and anemia (18–20). Consequently, many studies focus on single analytes or analyze blood only sparsely in time. A minimally disruptive alternative to blood draws is intravascular microdialysis, where a semipermeable membrane continuously samples the unbound fraction of bloodstream molecules below the molecular weight cutoff, while leaving platelets and other large molecules in circulation. After calibration, dialysate sample concentrations can be used to track changes in plasma levels (21–26), resulting in measurements that are not significantly different from blood draws for several key molecules (27). Clinically, both interstitial and intravascular microdialysis have already enabled minute-by-minute lactate and glucose monitoring (28) and steroid hormone profiling over 24hrs periods in humans (29). In rodents, simultaneous blood and brain microdialysis has been shown to be compatible with behavioral studies (25, 30). However, access to prolonged-use, easily accessible, and stable jugular microdialysis implants for mice is still challenging.

High-frequency blood dialysate samples can be analyzed with modern Liquid Chromatography-Mass Spectrometry (LC-MS) workflows, such as differential chemical isotope labeling (CIL), to measure hundreds of bloodborne compounds with high sensitivity (26, 31–33). The multiplexed nature of the measures enables high-dimensional (high-D) analyses of hormonal data. So far, both experimental and theoretical work have been constrained to single or low-dimensional (low-D) endocrine control motifs, such as glucose-insulin-glucagon (34, 35), or HPA cortisol control (36–38). We are inspired by the success of state-space methods applied to large-scale neural data, gene regulatory networks, and metabolomic profiles, which have demonstrated that high-D biological data can collapse onto low-D manifolds (32, 39–43). Given that neural activity, gene regulatory networks, and bloodborne messengers have shared dynamical features, such as coupled feedback, varying timescales, and coordinated endocrine axes, we wondered if bloodborne amines and small peptides dynamics also admit a low-rank physiological state space, and whether those trajectories align with behavioral axes.

Here, we introduce a workflow that incorporates chronic jugular microdialysis implants in freely moving mice, paired with chemical isotope-labeling LC-MS analyses of amine metabolites and small peptides for state-space reconstruction. We demonstrate the ability to measure hundreds of bloodborne compounds with stable quality control over nearly 8 hours at a 7.5-minute cadence. This data is compatible with a high-D, low rank description, with the first 10 principal components explaining more than 70% of the variance. Furthermore, low-D projections significantly correlate at different delays with locomotion of the animal, opening the door to systems-level physiology-behavioral studies.

## Results

### Chronic microdialysis (MD) jugular implants in freely moving mice enable multiplexed, frequent measurements of plasma compounds

Microdialysis (MD) is a technique that uses a semipermeable membrane to collect dialysate samples from cerebrospinal fluid (CSF), blood, or tissue (44). Most MD applications are acute, or performed within 1-3 days of probe implant (25, 30, 45). Chronic MD probe implants in rats have been used in the brain to monitor circadian rhythms and in the blood to measure drug concentrations, remaining patent for more than 1 week and up to 4 weeks (27, 46). In mice, tissue and vascular MD applications have been shown to be feasible in anesthetized and awake mice (25, 32, 47, 48). However, unlike whole-blood catheters (19), chronic MD jugular implants that remain patent for at least 7 days and are easily accessible *ad libitum* have, to the best of our knowledge, not been employed in freely moving animals. This is likely due to classical MD probes using a bulky liquid cross and stiff materials (Supp. Fig. S1), which make implantation harder than standard jugular catheters and hinder recovery and movement.

To overcome this, we designed an optimized probe and a surgery with key enhancements. First, we employed probes with a smaller liquid cross and suture point, enabling chronic implantation, and funnel-shaped flared tube endings and soft tubing materials (Fig. 1A). This allows the probe to be implanted more easily, while also allowing the animal to move more comfortably. The flared tube endings are designed to be connected to a dual-channel vascular access button (VAB), which is sutured between the scapulae of the animal. During surgery, the probe membrane is implanted into the jugular and advanced cranially with standardized placement (Fig. 1B; Supp. Fig. S2). The tubes loop around the chest through a generous tunnel (Fig. 1C, Supp. Fig. S2). This creates a system that is accessible *ad libitum* by connecting the VAB to a flow pump and a fraction collector for dialysate samples collection (Fig. 1C-D), and also for delivering perturbations (21). The full setup consists of an enclosure with a dual-channel swivel that allows the animal to move freely (Fig. 1D).

**Figure 1:**
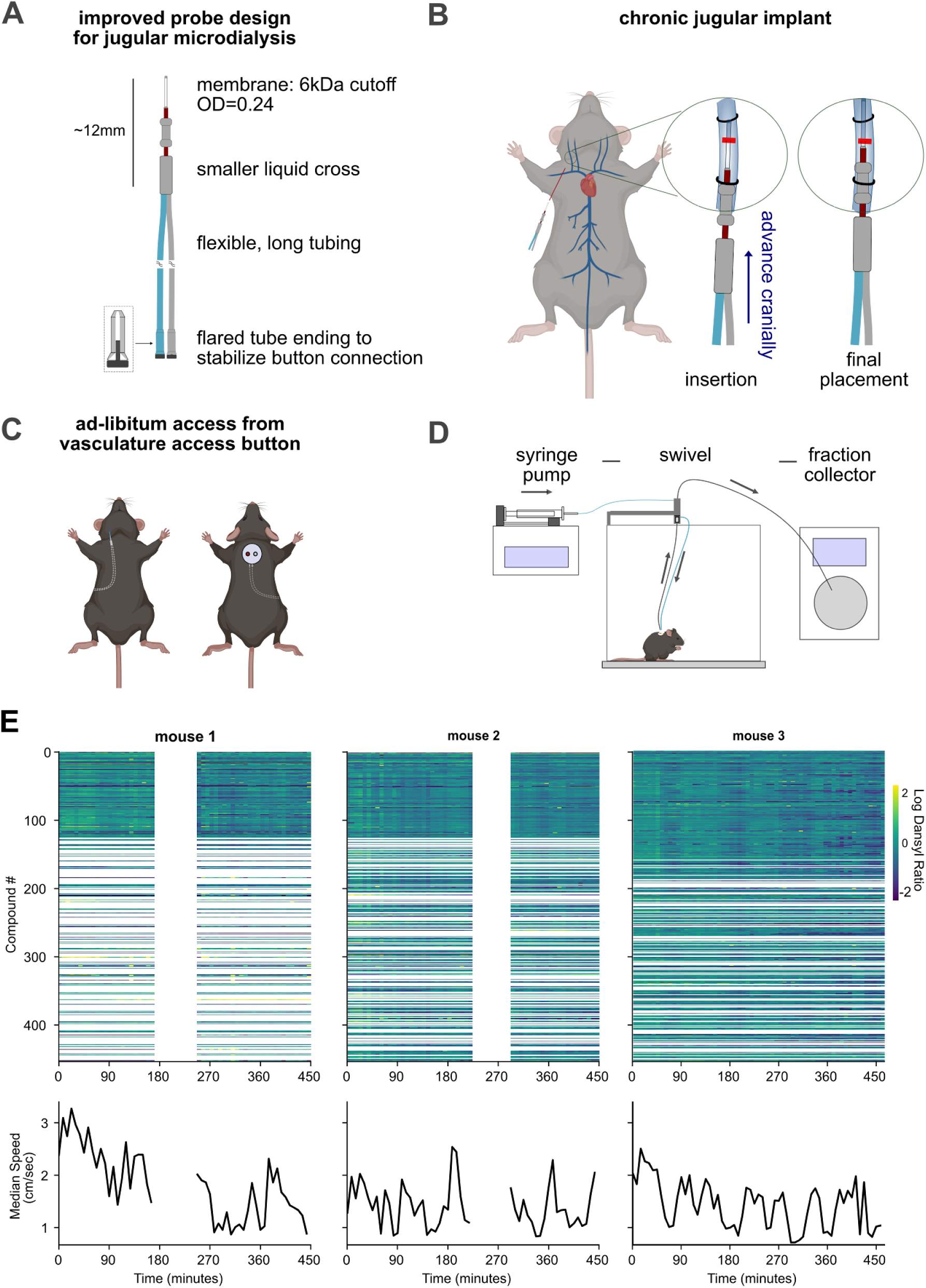
Chronic jugular microdialysis enables high-frequency, multiplexed blood chemistry in freely moving mice. **(A)** Microdialysis (MD) probe with optimized design for chronic jugular implants. Inlet line (blue) and outlet line (gray) connect to the vascular access button (VAB), transition through the redesigned liquid cross, and continue into the probe shaft (red), terminating in a 3.0-mm-long semi-permeable membrane optimized for vascular MD with 6 kDa molecular weight cutoff. **(B)** Surgery cartoon showing the key step for cranial probe advancement and suturing (details in Supp. Fig. S2) **(C)** The flexible tubes loop around the chest with ample space to avoid strain and promote recovery, making the MD implant easily accessible for at least 7 days. **(D)** Experimental flow: syringe pump continuously pumps a saline solution through the jugular MD probe via swivel/tether, which is then collected into a refrigerated fraction collector. Samples are then analyzed offline using differential chemical isotope labeling liquid chromatography-mass spectrometry **(E)** Representative 6-8 h recordings (7.5-min sampling) show hundreds of validated, high-quality dialysate compounds alongside average locomotion (7.5 min intervals aligned to MD samples) in three mice, with implants patent for >7 days post-implant. Common 123 high-quality compounds detected across all three animals are shown at the top. White gaps represent pauses in the recordings or compounds not detected in all animals.

Using an integrated microdialysis, ultra-high performance liquid chromatography, and mass spec workflow with differential chemical isotope labeling, we obtained stable, time-resolved profiles of bloodborne amines, amino acids, small peptides, aromatics, and other molecules (Supp. Fig. S3; Supp. Table 2). In two animals, a parallel aliquot enabled targeted quantification of steroid hormones, with integrations performed in Skyline (Methods; data not shown), while we performed untargeted amine metabolomics in all three animals in IsoMS Pro 1.2.7. Across runs, a background ion at m/z 251.0849 was consistently observed and used to verify mass accuracy. Its presence and mass, invariant across runs, together with all scan m/z values falling within expected ranges, indicate stable acquisition and mass accuracy throughout the dataset (Supp. Fig. S3; Methods).

Across the 3 animals, we detected 1069, 1471, and 1376 compounds, respectively. Out of those, 241, 343, and 345 passed stringent criteria for RT with an error below 60 seconds (Methods). The intersection across the three animals, including further restrictions in terms of consistency and replicability across technical replicates (Supp. Fig. S3), consisted of 123 compounds (Methods). A Bland-Altman plot showed no systematic bias for high or low chemical isotope labeling ratios (Supp. Fig. S3D).

With this setup, we monitored the free fraction of hundreds of amines and small peptides at 7.5-minute cadence for 6-8 hrs in 3 freely moving mice, together with the speed of the animal, calculated from keypoint tracking from videos recorded from an overhead camera (Fig. 1E). The implants maintained patency for at least 7 days post-implant in each animal. In one animal, we performed a control experiment to quantify the relative recovery of analytes at different flow rates. We found that a flow rate of 1.2 μL/min guarantees the optimal compromise between reliability and temporal precision of the measurement (Supp. Fig. S4). This setup and data, including a comprehensive metabolomic profiling on microdialyzed rodent blood samples collected in vivo, allow us to ask how physiological processes relate to behavior.

### Single-compound analyses recover known relationships between metabolites and behavior and reveal new ones

We measured the dynamics of hundreds of compounds and concomitant locomotion over 6-8 hours in three mice (Fig. 1E; Supp. Fig. S5). To obtain an overview of the relationship of single compounds and locomotion, we computed the cross-correlation for all 123 compounds that were validated across all three mice with locomotion (Supp. Fig. S6). Overall, we found that 87 compounds had significant (p<0.05, Bonferroni corrected) correlation with locomotion, out of which 74 had positive correlation, and 13 negative. Several relationships suspected in prior work emerged clearly here. To validate this acquisition method as a discovery tool, we looked specifically at compounds with known relationships between endocrinology, physiology, and metabolism with behavior (Fig. 2A).

**Figure 2:**
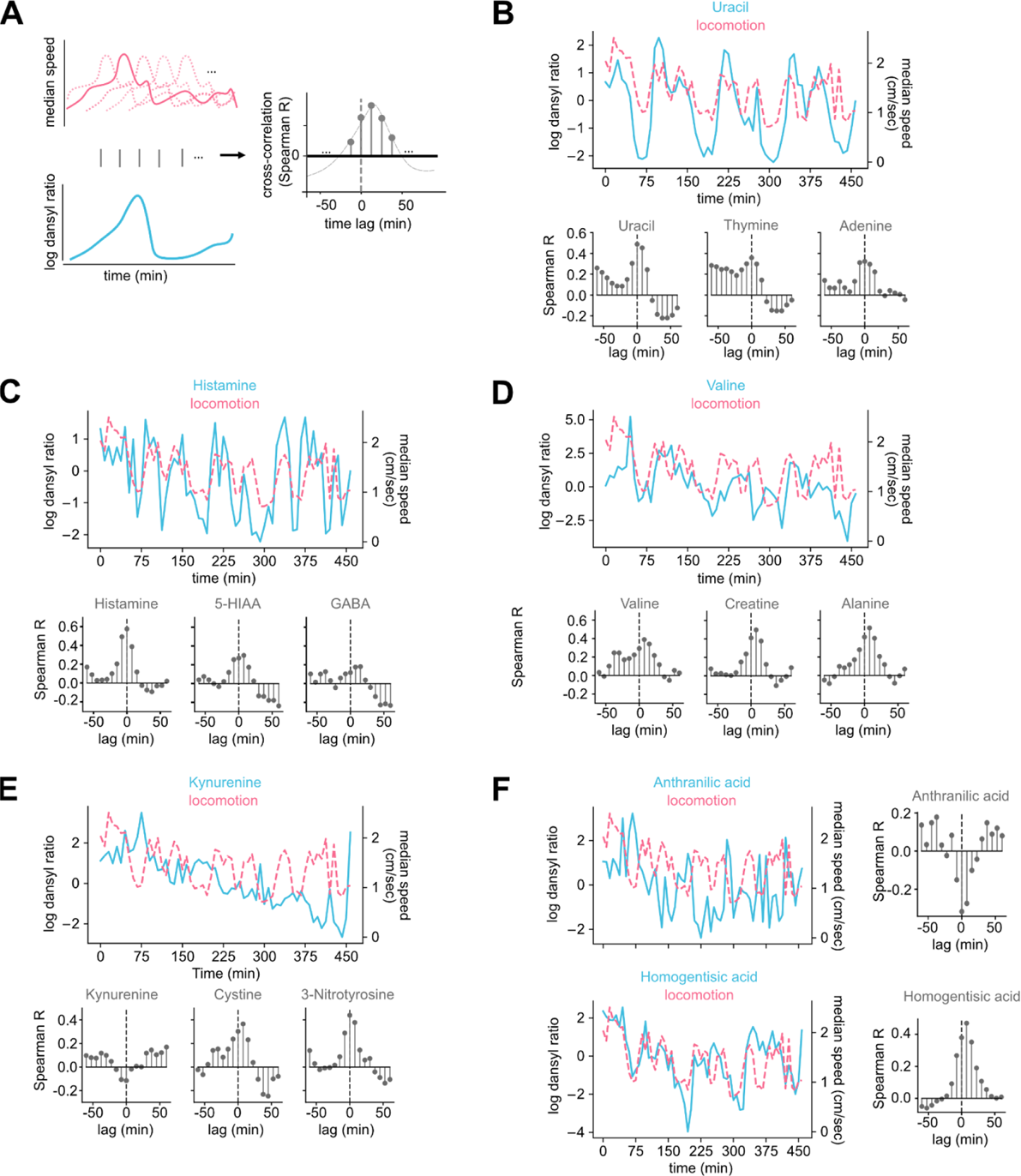
Single-compound dynamics recapitulate known physiology and reveal new behavior-linked relationships. **(A)** Cross-correlation workflow: locomotion time series is shifted over lags to quantify delayed alignment (Spearman r). **(B)** Purine turnover during movement: Uracil, Thymine, and Adenine correlate with locomotion at 0-7.5-min lag, consistent with exercise-driven pyramidine and purine catabolism. **(C)** Locomotion-related biogenic amines and metabolites: Histamine peaks near zero lag; 5-HIAA (a serotonin turnover marker) shows delayed correlation (∼7.5 min); GABA exhibits a brief onset-aligned change followed by a reduction. **(D)** Amino-acid and energetic pathways: Valine (BCAA), Alanine, and Creatine show positive correlations peaking ∼0-15 min after locomotion, consistent with nitrogen handling, Cahill cycle, and phosphocreatine replenishment. **(E)** Stress/redox signatures: Kynurenine displays negative association with locomotion; Cystine and 3-Nitrotyrosine show positive relationships. **(F)** Additional associations not classically linked to locomotion but biologically plausible via pathway context: Anthranilic acid and Homogentisic acid. All locomotion/compound time series examples in this figure derived from mouse 3, while cross-correlograms were computed on data from the three animals combined.

For example, it is well known that muscle contraction and sympathetic arousal ramp up ATP turnover, leading to an increase of purine and pyrimidine catabolites within minutes (2, 49–51). To that extent, we measured the Spearman cross-correlation of adenine, uracil, thymine, (Fig. 2B) and xanthine (Supp. Fig. S6) with locomotion, and found a positive correlation within 0∼7.5 minutes lag. This confirms that our approach captured exercise-induced purine and pyrimidine turnover, which is a known physiological benchmark that, to our knowledge, had not previously been resolved with this temporal resolution in freely moving mice.

Next, we examined compounds related to exercise (Fig. 2C). We analyzed 5-HIAA, which acts as a readout of serotonin turnover/metabolism (52) and is known to rise after aerobic bouts (53). We also evaluated histamine, because exercise triggers mast cell histamine formation and post-exercise vasodilation (54, 55). Finally, we looked at GABA, the chief inhibitory neurotransmitter in the brain (56). We found a high correlation between microdialysis-measured histamine and locomotion at ∼0 lag, and a delayed agreement of 5-HIAA with locomotion at a lag of 7.5 minutes, as expected from the literature (57, 58). GABA exhibits a weak peak aligned with the onset of locomotion and a drop after ∼22.5 to 45 minutes. The relationships we observe here highlight the potential of chronic MD to track peripheral correlates of exercise, and could be used as an exploratory tool to find correlations between states and behavior (59, 60).

We next measured the relationship between various amino acids and locomotion (Fig. 2D). Working muscles exchange amino acids with plasma through the ammonia detox and the glucose-alanine cycle, also known as the Cahill cycle (49). Branched-chain amino acids are oxidized in increased amounts during exercise and need to be replenished afterwards (61, 62). Accordingly, we found valine levels to be highest with a 7.5 minute delay compared to locomotion. Alanine is expected to increase with locomotion as part of the glucose-alanine cycle, where muscle cells release alanine to the liver for gluconeogenesis (63). We found that alanine was strongly modulated by locomotion at a short delay, in agreement with the literature. Finally, we analyzed creatine levels compared to locomotion, and found the strongest overall correlation at a delay of 7.5 minutes. This is consistent with creatine’s role in replenishing phosphocreatine stores in muscle (64). This served as an internal validation, showing that these data captured muscle energy flux with the expected delay kinetics. The ability to track coordinated amino acid and creatine dynamics over minutes provides a resolution not achievable with previous bulk blood sampling approaches.

We also analyzed metabolites in the kynurenine pathway and other markers of locomotion-induced redox/stress (Fig. 2E). To that extent, we measured the cross-correlation of kynurenine, cystine, and 3-nitrotyrosine with locomotion. Kynurenine shows a negative correlation with locomotion at zero lag, a result expected from the literature, as kynurenine and its downstream byproducts were found to be altered in older adults during and after exercise (65). Cysteine shows an interesting trend, with a strong positive correlation at 7.5 minutes lag, and a drop after 22.5 minutes. This might be due to cysteine’s roles in multiple pathways, including protein synthesis, antioxidant activity, and immune function (66). Finally, we analyzed 3-nitrotyrosine (3-NT), a biomarker of oxidative and nitrosative stress associated with various disease states, aging, and inflammation (67). We found 3-NT shows a strong positive correlation at no lag with locomotion. These compound-specific delays and polarity shifts suggest that chronic MD could be used to resolve distinct phases of oxidative and immune-related metabolism following activity, which is typically collapsed in single-endpoint blood assays.

Further, we found a negative correlation between anthranilic acid and locomotion (Fig. 2F). Anthranilic acid does not have a direct, established role in mouse locomotion; however, it is a metabolite of the kynurenine pathway, making the relationship we observed biologically plausible (68, 69). Finally, we also found an increase of homogentisic acid at 7.5 minutes after locomotion (Fig. 2F). Homogentisic acid is a metabolic intermediate, normally formed during the catabolism of phenylalanine and tyrosine, and is known for its involvement in Alkaptonuria, a rare genetic disease which leads to ochronosis, osteoarthritis, and loss of locomotory capability (70). Although consistent with its role in tyrosine breakdown and dopamine formation, this link to locomotion has not been previously reported. These findings exemplify how high-frequency, multi-compound monitoring can reveal transient interactions that could be missed in snapshot assays.

Finally, we looked for known compound-compound interactions, such as histidine and histamine, which is produced from histidine (71); arginine and citrulline, which can be produced by nitric oxide synthetase conversion from arginine (72); and kynurenine and N-formylkynurenine, which is produced in the same enzymatic pathway (73) (Supp. Fig. S8). Each pair, as expected, showed a peak correlation at 0 lag.

In addition to instantaneous behavioral correlations, we used the binned behavior data and looked for consecutive collection timepoints where animals had an overall lower or higher median speed. During rest, the body enters a state of fasting and repair (74). We compared median z-scored compound values during prolonged immobility vs. persistent locomotion, and found a clear negative correlation, indicating that many compounds are inversely modulated by locomotion (Supp. Fig. S7). The compound with the largest locomotion-related change compared to immobility was histamine, which plays a well-known role in wakefulness (75). Conversely, hydroxycinnamic acid, a microbial metabolite of aromatic amino acids, was most enriched during rest, as expected from literature on aromatic metabolism in humans (76). Together, these state-dependent shifts demonstrate that chronic MD can capture systemic biochemical signatures of sustained activity, which could be used to complement electrophysiological or imaging-based approaches.

Overall, these results demonstrate that chronic MD not only recovers established metabolic signatures of activity but also uncovers new, biologically plausible relationships that could reveal how behavior dynamically coordinates metabolism across multiple physiological systems.

### Physiological manifold with behavior alignment

Previously, we showed that analyzing single compound dynamics recapitulated known relationships with locomotion and also uncovered putative novel relationships (Fig. 2, Supp. Fig. S6-S7). Here we take a systems-level perspective to study the high-D structure of the data and the covariation across compounds (Fig. 3A). We asked whether blood messenger dynamics can be summarized by a small number of latent dimensions, and whether the projection of the data along these axes aligns with interpretable behavioral variables. To address this, we employed nonparametric, robust methods to estimate compound covariation (Spearman’s rank correlation) and standard linear decomposition (principal component analysis, PCA) to extract projections that capture the dominant modes of variance.

**Figure 3:**
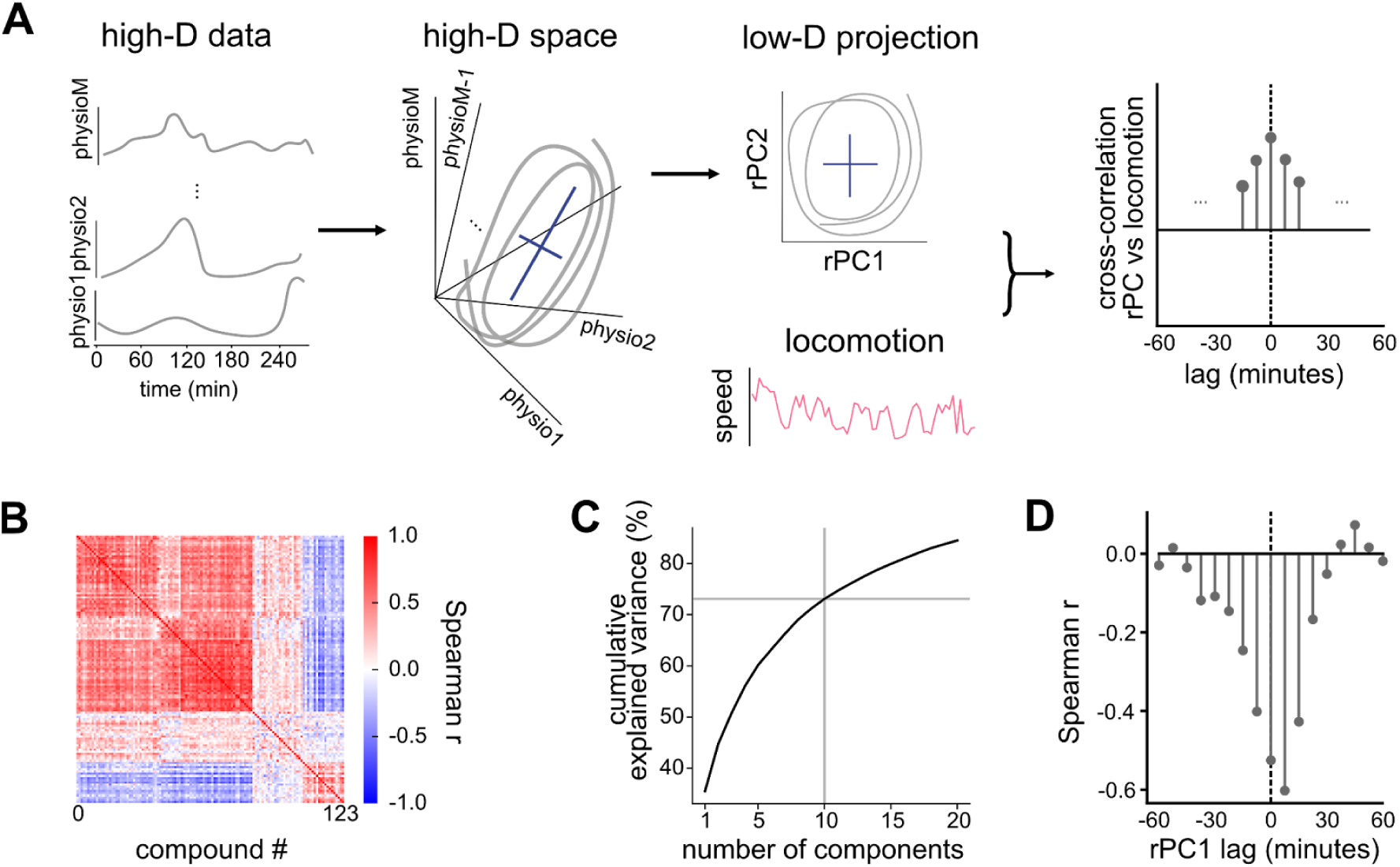
A low-dimensional physiological manifold aligns with behavior. **(A)** Analysis overview: high-D compound time series → covariance structure → low-D projections → behavioral alignment tests. **(B)** Spearman correlation matrix on combined datasets (Ward linkage ordering) shows prominent block structure (within vs. outside ratio = 4.17) **(C)** Robust PCA (rank based / Spearman) on the combined dataset: variance explained vs number of components shows ∼73.08% captured by 10 components, rPC1 explains 35.5%. **(D)** Cross-correlation between rPCs and locomotion shows rPC1 peaking at 7.5-min lag (gray dashed line marks zero lag). See Supp. Fig. S10 for Cross-correlations with other rPCs.

We begin by considering each compound as a time series: evaluating how it changes over time, and how it correlates with all other compounds. The resulting matrix exhibited rich structure, with hierarchical clustering (Ward linkage) ordering compounds into coherent modules (Fig. 3B). As a control, we confirmed that the Spearman correlation structure was highly similar across the three animals (all cosine similarities > 0.5; p<1e-10; Supp. Fig. S9A). Because hormonal dynamics may be influenced by locomotion, we also computed the correlation across compound timeseries discounting the effect of locomotion and observed similarly high correspondence across datasets (all cosine similarities > 0.5; p<1e-10; Supp. Fig. S9B). Finally, the Spearman correlation matrices from original and column-normalized data remained highly correlated (all r>0.8, p<1e-10, Supp. Fig. S9C). Together, these controls confirm that the correlation structure is stable across animals and robust to technical variation, providing a reliable foundation for analyzing the high-dimensional organization of physiological dynamics.

To examine the high-D structure of the data, we computed a Spearman-based, robust PCA, which is less sensitive to outliers and non-linearity/gaussianity than normal PCA, across the entire dataset of the three animals combined. We plotted the variance explained by each robust principal component (rPC) (Fig. 3C). We noticed that the first 10 rPCs explain 73.08% of the variance, with a gentle decay as we add dimensions. In particular, we notice that the first rPC explains 35.5% of the variance, and the successive rPCs contribute 5∼10% each. This indicates that the complexity of circulating molecular dynamics can be represented by a compact, low-dimensional manifold. This result is reminiscent of NMR-based single snapshot metabolomics across humans (41), but to our knowledge, it was not previously demonstrated for *in vivo, dynamic* metabolomic profiles recorded in freely moving animals.

Next, we examined whether projections along these rPC axes were aligned with locomotion. Cross-correlation analyses between each rPC and locomotion (Supp. Fig. S10) revealed that rPC1 was strongly correlated with movement (Fig. 3D, Spearman r=-0.59, p<1e-19), peaking at a 7.5 min lag. This suggests that rPC1 represents a coordinated physiological mode driven by locomotor activity. Inspection of rPC1 loadings showed the highest weights include many amino acids and small peptides: threonine, proline, alanine, glutamyl-methionine, serine, asparagine, glutamyl-alanine, and histidine. These are muscle- and nitrogen-related metabolites, and together this indicates the potential ability of our analyses to recover coherent groups of compounds whose collective dynamics track behaviorally relevant physiological processes. In conclusion, these findings reveal a low-dimensional physiological manifold in which behaviorally relevant factors, such as those linked to energy and amino-acid metabolism, emerge as dominant axes of variance. By bridging the single-compound and systems levels views, this approach helps uncover metabolic coordination with behavior that could not be inferred from traditional single-compound assays. The alignment between global physiological structure and behavioral dynamics highlights chronic microdialysis as a promising tool for discovering organizing principles of systemic and homeostatic regulation.

## Discussion

We present an experimental workflow that lays the foundation for reconstructing a latent, low-rank physiological manifold from high-frequency measurements of circulating molecules in freely moving mice. By combining chronic jugular microdialysis (MD) with chemical isotope labeling LC-MS, we track hundreds of amines and small peptides at 7.5-min resolution over hours (Fig. 1). Single compound dynamics recover known physiological relationships, such as purine turnover, locomotion-related biogenic amines, and amino acid metabolism (Fig. 2). Dimensionality reduction methods reveal a compact manifold whose trajectories align with behavior, such as locomotion (Fig. 3). Together, these results establish technical feasibility, biological validity, and a systems level description that shifts the focus from single, isolated hormones to coordinated trajectories in a latent physiological space.

### Why microdialysis and mass spectrometry?

Systems and behavioral neuroscience studies could benefit from interdisciplinary methods to study the drivers of behavior (77). This includes studying how physiological needs are transmitted to tissues and the brain, and how they influence behavior (78). Many existing readouts in metabolism and endocrinology studies are downstream byproducts, such as sweat (15, 79), urine (80), and feces (81, 82). These products integrate multiple upstream processes and can be delayed or confounded by local tissue kinetics; for example, fecal stress markers are delayed on average by 4∼12 hrs compared to plasma levels (83–85). Intravascular MD directly samples the bloodstream, capturing the unbound fraction of circulating messengers within the probe’s molecular-weight cutoff (21, 25), enabling continuous collection without repeated animal handling stress, and captures minute-scale dynamics (26). Moreover, given that there is no net fluid exchange, MD is compatible with hours-long, high-frequency sampling (21, 44). The most similar application is jugular catheter implant for blood draws in freely moving mice (19). Although this method permits fewer daily samples than microdialysis, it offers the advantage of providing a comprehensive measure of bound and unbound compound concentrations, as well as large-scale components, such as platelets and proteins (86).

Compared to enzymatic/electrochemical sensors, such as continuous glucose monitors, MD trades bandwidth (sub-minute intervals) for breadth (hundreds of analytes). With LC-MS, analysis of dialysate expands the chemical scope to a broad panel of amines and small peptides (31, 33, 82) with minimal cross-reactivity or sensor drift, at the cost of lower temporal bandwidth and temporal convolution. Moreover, LC-MS analyses can flexibly incorporate targeted panels (steroids (87), eicosanoids (88), or large peptides/proteins (89)) so long as they fall within recovery constraints.

Finally, the jugular vein provides a suitable sampling site due to its high flow and robust plasma access. Most cerebral venous blood drains into the external jugular veins, making it an ideal placement for sampling blood messengers related to neuroscience. The small membrane and shaft footprint, softer tubing, and compact liquid switch employed here were essential to making a chronic, behavior-compatible jugular implant feasible in mice, resolving the contradiction between probe bulk/stiffness and the small vasculature (Supp. Fig. S1-S2). Together, MD–MS offers a scalable approach to capture systemic dynamics across many biochemical axes simultaneously.

### Transition from single analyte analyses to a physiological manifold view

Our single compound analyses reproduce well-known relationships, such as movement-related purine turnover (50), histamine and 5-HIAA changes consistent with post-exercise vasodilation (54, 55), serotonin turnover (90), and amino acid activity related to energy and nitrogen handling (50). However, zooming out and focusing on a systems-level view, the covariation across hundreds of compounds uncovers interesting facts. For example, principal component analyses capture a large proportion of variance with few dimensions, with leading axes aligned with interpretable behavior (e.g., velocity vs. rPC1, Fig. 3D). This observation echoes the low-rank structure described in neural population activity (39) and gene regulatory networks (40), providing a reduction in complexity to a handful of latent variables. Low-rank structure identification has been a powerful tool in neuroscience, uncovering principles of motor control (91) and decision-making (91). These results suggest that endocrine and metabolic control may operate over a compact set of latent variables, contributing to a growing body of evidence that demonstrates these systems-level methods can be useful in metabolic research (32, 41–43).

### Future applications and extensions

In addition to sampling analytes, microdialysis also enables the delivery of specific molecules and pharmaceuticals (21). This paves the way for future open-loop and closed-loop experiments, where targeted perturbations can be performed at will; for example, when the hormonal manifold position falls within a certain area of state space or an animal performs a specific behavior, such as eating. Closed-loop perturbations tied to the real-time readout of hormonal levels (92) could include optogenetic stimulation of various brain regions (e.g., hypothalamus), pharmacological endocrinology manipulations, or nutrient infusion/delivery. On the modeling side, computational methods can be used to discover low-rank dynamics from high-dimensional data (94, 95) and detect endocrine-behavioral interactions over different timescales (93). This workflow can also be integrated with other physiological measurements, such as muscle EMG, implanted thermal RFID tags, or neural recordings, thus enabling bidirectional mapping between brain activity, effector systems, and circulating bloodstream messengers.

## Conclusion

Chronic jugular MD in freely moving mice, paired with isotope-informed LC-MS, reveals that complex physiological dynamics collapse onto a low-dimensional manifold with its main axis aligned with locomotion. This work provides both a technical foundation for minimally disruptive, high-throughput monitoring and a conceptual shift toward a dynamical-systems physiology of homeostasis.

## Methods

### Surgery

All surgeries were done in accordance with animal protocols 25-0281 and 24-0263 approved by the Janelia Research Campus’s Institutional Animal Care and Use Committee. All measurements were performed on single-housed C57BL/6CRL mice between the ages of 11-15 weeks and between the weights of 26-35g. Surgeries used sterilized supplies and aseptic techniques in a dedicated surgery space and a stereoscope. Mice were anesthetized with 3% Isoflurane to reach a surgical plane as confirmed by toe pinch. Following induction, the ventral neck (mandible to clavicle) and dorsal thorax (neck to rib cage) were shaved, then cleansed with 70% ethanol. Animals were transferred to a surgical platform, modified to allow passage of the vascular access button (VAB) through a central opening. The platform was prepared with a nosecone and a heat source on a sterile field prepared with Press’N Seal film. Surgeons applied ophthalmic lubricant, provided Ethiqa XR (3.25mg/kg), and surgical sites were scrubbed sequentially with ethanol and chlorhexidine. Animals were sterilely draped with Press’N Seal film wrapped around the torso to retain body heat and act as a sterile drape. Surgeons applied marcaine to the incision sites (maximum total dose 7mg/kg). A 6-9mm dorsal incision was made between the scapulae, and a subcutaneous pocket was blunt-dissected for VAB placement. Rotating the animal as needed, a channel was created subcutaneously from the dorsal incision to the ventral neck. In the supine position, a 6-9 mm ventral incision was made lateral to the midline over the right jugular vein (clavicle to mandibular ramus). The VAB was connected to the microdialysis probe, and tubing was tunneled subcutaneously from the dorsal incision to the jugular site, keeping the tip of the probe capped. The dorsal incision was closed around the VAB with vicryl suture using a subcuticular continuous suture pattern. The VAB was connected to a CMA 4004 Syringe Pump with a T1 isotonic sterile perfusion fluid for peripheral tissue at 10 μL/min for 10 minutes, and then reduced to 0.3 μL/min for the remainder of the surgery. The jugular vein was exposed by blunt dissection. Approximately 5-7 mm of the vein was isolated, and two 6-0 silk ligatures were passed beneath the vessel; one placed caudally at the widest portion of the vein, and the other cranially, immediately caudal to a small landmark tributary vein, spaced ∼3-4 mm apart. Knots were tied in each silk suture but kept loose enough as not to occlude the vessel. The probe was fed through the loose caudal suture exterior to the vein and aligned to be inserted. Hemostats were tightened on the tails of the suture to partially occlude the vessel and reduce bleeding. A small transverse incision was made in the vein between the two ligatures, and the probe was gently inserted into the vessel lumen and advanced cranially past the cranial ligature. The cranial ligature was tightened to secure the probe within the vessel, followed by the caudal ligature. Proper functioning of the probe tip was confirmed by noting the constant presence of dialysate in the tube exiting the VAB. The skin incision over the vein was closed with Vicryl suture using a continuous subcuticular suture pattern. A protective metal cap was placed over the VAB to prevent damage. Post-operative analgesics (ketoprofen 5mg/kg or meloxicam 2mg/kg) were administered daily per protocol, and animals were allowed to recover under observation.

### Probe design, membrane cutoff, perfusate, and flow rate

Probe design followed a design as in Fig. 1A and Supp. Fig. S1B. The membrane molecular weight cutoff was 6kDa. We employed CMA probes with a smaller liquid cross and suture point, funnel-shaped flared tube ending for easy yet stable connections, and softer tubing materials for ease of probe implantation and animal movement. Microdialysis experiments were conducted 5 to 10 days after implantation of the probe. he During experimental measurements, probes were connected to a syringe pump, and perfused with saline solution at a flow rate of 1.2 μL/min. Microdialysis samples from the blood were collected every 7.5 min. The collected samples were kept at 6-8°C during the sampling and transferred to -80 °C immediately after the end of the experiment.

### Experimental equipment and procedures

#### Sample collection during behavior

Behavioral sampling was conducted in a small, circular arena (20cm diameter) equipped with an overhead camera (Logitech C920). The microdialysis behavioral setup consisted of:

- Syringe pump (CMA 4004)
- Dual-channel liquid swivel (Instech 375/D/25) mounted over a CMA mouse cage
- Refrigerated fraction collector (CMA 470)

The collection procedure consisted of: 1) Pre-recording (setup and stabilization): load and prime syringe, start devices, flush the probe, and verify stable flow through the swivel and collection lines; 2) Recording (behavior + sampling): Tether the animal via the VAB and swivel and record behavior while collecting dialysate into time-stamped vials; 3) Post-recording shutdown and cleanup.

#### Routine probe maintenance (Periodic flushing)

To maintain patency and minimize tissue buildup around the vascular access, the microdialysis probe was flushed once daily. During the first 2-3 postoperative days, depending on the animal’s condition, wound healing, and overall recovery, flushes were performed under general anesthesia to reduce tissue strain (jugular/muscle/skin). Thereafter, daily flushes were conducted in awake animals.

#### Computing transit delay and empirical validation

Transit delay refers to the time it takes for the liquid to travel from the tip of the probe, where it begins to interact with the blood, to the fraction collector. To do that, we need to add: half probe DV + VAB button DV (one connector) + tube from VAB to swivel + swivel center channel + tube from swivel to fraction collector. For tubing with an inner diameter (ID) 0.12 mm (radius *r* =0.06 mm), the segment volume is: *V* = *l* × π × 0.06^2^ where *l* is the length in mm.In our experiments, we calculated an 18.6 μL dead volume, including the swivel, VAB, connectors, and probe. This, divided by 1.2 μL/min, yields approximately a 15-minute delay (two 7.5-minute “time bins”), which needs to be accounted for in the analysis. We measured this also in the lab, where instead of a probe we connected the swivel with a 25cm tube length (which is similar to the dead volume of MD probe + VAB), and measured 14min 38sec. This is compatible with our theoretical measure, and we considered 2 time windows (15 minutes) as the delay.

#### Dilysate analyses: dansylation and isotope dilution LC-MS/MS + pooled-quality control and internal standard library

We paired the microdialysis sample collection with ultra-high performance liquid chromatography (UHPLC)-mass spectrometry (nanoLC-MS) and differential chemical isotope labeling workflow specifically designed to measure bloodborne amines and phenols (Supp. Fig. S3). One aliquot of the dialysates was subjected to nanoLC-PRM (Parallel Reaction Monitoring) for steroid hormone analysis using an Orbitrap Ascend Tribrid mass spectrometer equipped with a Vanquish Neo UHPLC. The PRM data were analyzed using Skyline (V24.1). Global metabolomic analysis was done through Compound Discoverer 3.3. Chemical isotope labeling was performed on 5 µL of the dialysates at each time interval using the dansyl-labeling Kit from NOVA MT and then analyzed using a VanquishTM Horizon UHPLC connected with an Orbitrap Fusion Lumos Tribrid mass spectrometer (Thermo Fisher Scientific, Bremen, Germany). M/z 251.0849 was selected as a background peak to check the mass accuracy for the samples. The accurate mass and isotope pattern of the peak were consistent during all runs, and all scanned m/z are within the expected tolerance, showing good stability and mass accuracy for the data acquisition.

#### LCMS data processing and selection of compounds

The resulting data was interpreted using software IsoMS Pro 1.2.7 (Nova Medical Testing), to extract peak pairs and calculate the intensity ratio of each peak pair. For metabolite identification, based on multiple metabolite identifiers, peak pairs were searched against the chemical isotope labeling labeled metabolite library and linked identity library.

Furthermore, after IsoMS Pro software processing (94), we applied the following steps for data cleaning and compounds selection:

1. <u>Selection of high-quality compounds:</u> Based on IsoMS Pro 1.2.7 (Nova Medical Testing) data readout, we selected compounds with Tier 1 and Tier 2 detection quality (RT error < 60sec and reliable compound detection).
2. <u>Fraction outlier detection:</u> remove samples with median ratios too low (below median - 4×MAD)
3. <u>Technical replicates averaging:</u> in log2 scale, we computed the absolute difference among replicates, and set to blank if |rep1−rep2| > 1. We then averaged remaining replicates to produce one value per timepoint.
4. <u>Selecting consistent subset across animals:</u> We removed compounds from further consideration if they were missing more than 25% of the LCMS time points for any individual animal. Intersection set consists of compounds passing the criteria in all three animals.
5. <u>Interpolation of missing values:</u> each compound time series was linearly interpolated using scipy.interpolate.interp1d.
6. <u>Z-scoring for manifold analyses:</u> data of each compound is centered and scaled within each animal using robust z-score [(value−median)/MAD], and then data was concatenated for pooled analyses.

#### Behavioral data acquisition and processing

We acquired videos using OBS Studio and a Logitech C920 webcam, acquiring video at 30Hz and 1280 x 720 resolution. For all mice, neck position was tracked using DeepLabCut 3.0.0rc13 (Mathis et al 2018, Lauer et al 2022) with ResNet-50 convolutional neural networks (He et al 2016). The position and lighting of the arena changed sufficiently between animals to require a separate network for accurate position identification. The first two mice were recorded under identical conditions, and we trained the network with 51 manually labeled frames, and tested on 9 held out frames. On the test set, the model achieved RMSE of 7.56 pixels (0.40 cm; 2.0% of the 20 cm arena diameter), with mAP of 80.85% and mAR of 81.11%. Training performance (RMSE: 2.23 pixels, mAP: 98.92%, mAR: 99.22%) showed a train/test ratio of 3.4x. This network was applied to the four consistently recorded videos. The second network was trained on 80 training frames and tested on 4 test frames. The network achieved a test RMSE of 5.24 pixels (0.28 cm), with mAP of 92.71% and mAR of 94%.

Behavioral videos were analyzed at 30 fps using their respective trained networks. For each video, the x,y tracking coordinates were converted from pixels to centimeters using video-specific calibration measurements. A representative frame was obtained from each video, and the circular arena boundary (known diameter: 20 cm) was traced over the image using Adobe Illustrator’s Ellipse tool, and the pixel-to-centimeter conversion factor was calculated from the measured pixel diameter. Calibration factors averaged 0.0530 ± 0.0006 cm/pixel (mean ± SD) across all videos.

We first removed outliers. We considered two types of outliers: those that had x,y positions outside of the arena boundary, and any change in coordinates not biologically reasonable. For example, occasionally a mouse’s reflection placed the animal’s x,y position outside of the arena, which would be replaced with NaN (2,111 of 2.4 million frames). Similarly, any single-frame jump in coordinates 10 times higher than the 99th percentile of observed speed in that video was replaced with NaN (97.5-142.0 cm/s, 1,079 of 2.4 million frames).Using these filtered data, we calculated a 5-frame centered rolling median filter was applied to smooth noisy coordinates and camera sensor jitter. The filter computes the median from available values within each 5-frame window, treating flagged outliers as missing data points that do not influence the smoothed trajectory. The 5-frame window (0.167 seconds at 30 fps) simultaneously allows for the removal of high-frequency noise while preserving the mouse’s biological movements. We computed the speed of the animal as the Euclidean distance between consecutive coordinates, divided by the time between frames.

#### Statistics and Multivariate Data Analyses

Dialysate values are treated as arbitrary units. All analyses focus on within-analyte dynamics, using per-animal, per-analyte normalization (robust z-score using median and interquartile range). We do not interpret absolute extracellular concentrations nor between-analyte amplitude differences. We assume constant recovery over time for each animal and each given analyte. This implies that the measurements are scaled by a constant factor, hence Z-scoring per compound removes that scale, leaving dynamical properties, such as those measured by cross-correlation, unaffected.

#### Regression analyses

Linear regression was used for exploratory relationships between compound concentrations and behavioral covariates (e.g., locomotion velocity). Prior to regression, compounds were z-scored within an animal to remove baseline offsets. When applicable, variables were mean-centered to ensure the interpretability of intercepts.

#### Spearman cross-correlation

To assess temporal relationships between compound abundance and locomotion, we computed Spearman rank cross-correlations between each compound time series and the locomotion trace, using non-parametric rank correlation to reduce sensitivity to outliers. Cross-correlations were computed over symmetric time lags (−60 to +60 min) using a sliding-window implementation in scipy.stats.spearmanr, and reported as the maximal absolute correlation coefficient within the lag range. Peaks and their lag positions were used to infer leading or delayed relationships relative to behavior.

#### Partial correlations

To quantify correlations among compounds independent of locomotor influence, we computed partial Spearman correlations. For each pair of compounds x, y, and confound variable z (locomotion), the partial correlation was computed as the residual correlation between the rank-transformed residuals of x and y after regressing out z.

#### Immobility vs locomotion analyses

We selected two thresholds: locomotion was defined as med + MAD/2 (med = median of all animal’s medians, MAD = median absolute deviation), and immobility was defined as below med - MAD/2 for at least 2 time bins. We then analyzed the average z-scored value of each compound during periods of immobility vs locomotion.

#### Robust PCA

We performed dimensionality reduction using a Spearman-based robust PCA to capture low-rank structure while minimizing influence from outliers and non-Gaussian distributions. Each compound time series was rank-transformed, concatenated across animals after z-scoring, and decomposed using the robust covariance estimator implemented in sklearn.decomposition.PCA with Huber loss. Components (rPCs) were ordered by explained variance. For behavioral alignment analyses, each rPC time course was cross-correlated with locomotion as described above. Absolute loading values were used to identify compound groups contributing to each latent axis.

## Code availability

Original scripts used to generate all the results in this paper, including preprocessing and supplementary figures, are available at: https://github.com/michnard/Microdialysis_Paper All analyses were performed in Python (v3.9 and 3.14) using numpy, pandas, statsmodels, and scikit-learn packages, with detailed package and environment specifications present in the file “environment.yml” on the GitHub page for full reproducibility.

## Data availability

Original LCMS data, as well as preprocessed behavior tracking data, were deposited to FigShare: https://doi.org/10.25378/janelia.30556511

## Authors Contributions

MN: Michele Nardin, NW: Nan Wang, SE: Soad Elziny, CB: Claire Boyer, VP: Vojko Pjanovic, LS: Luisa Schuster, PB:Peter Boklund, SL: Sarah Lindo, KM:Kendra Morris, AI:Anoj Ilanges, JV: Jakob Voigts, EJD: Emily Jane Dennis

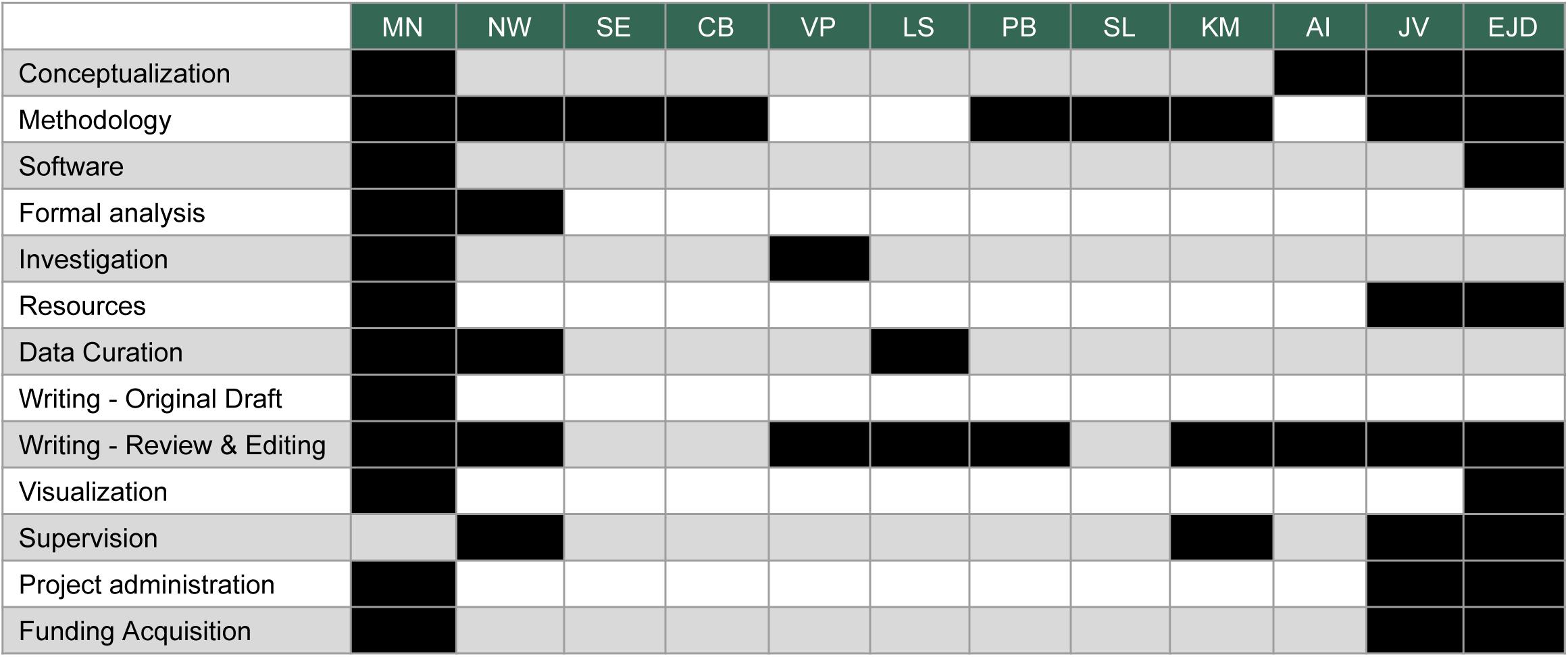

## Acknowledgements

The authors wish to thank Paul Tillberg, Tosif Ahamed, Virginie Ruetten, Pavlo Bulanchuk, Magdalena Schneider, and Ahmed El Hady for discussions, ideas, and support; Guinevere Bell for discussions about surgery and materials; Anne Kuszpit and the whole surgery team for help with surgeries and brainstorming; Rachel Gattoni for Vivarium support; Henrik Jessen for discussions about probe materials and development; Paul Loughnane for setup guidance and discussions on hardware and consumables; Megan Zipperer for help and suggestions on writing style and readability; Boaz Mohar for suggestions on figure schematics and exposition; Wei Wu for assistance in Mass Spec sample prep and analyses. MN used Grammarly for grammar check and Biorender for schematics. Link to license: https://BioRender.com/lxgr5j6

## Supplementary Material

### Supplementary Figures

**Supplementary Figure S1:**
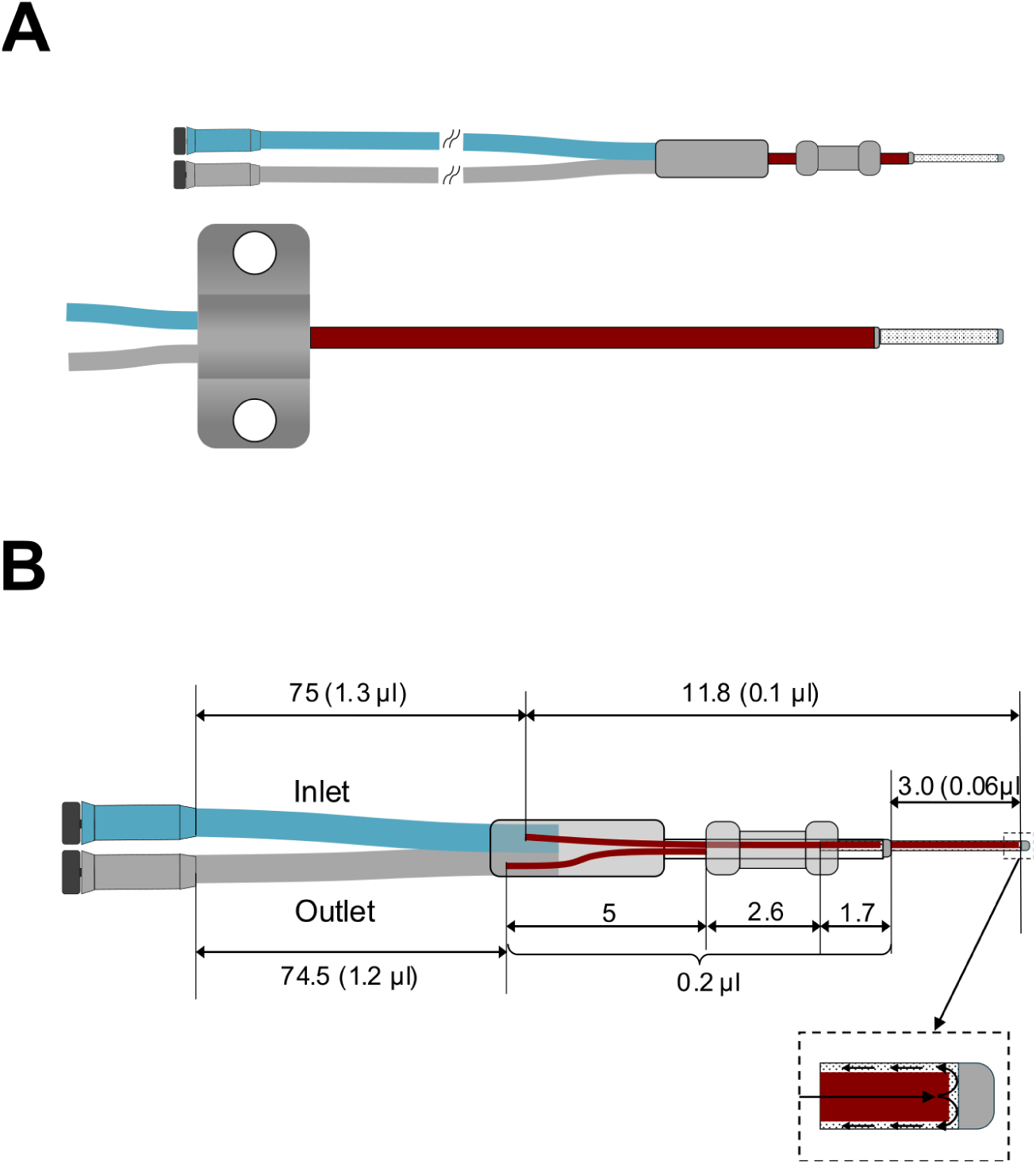
Probe design details. **(A)** Comparison of our optimized, redesigned MD probe (top) vs standard MD probe for tissue sampling (to scale). **(B)** Dimensional schematic and internal volumes: Lengths are in mm, dead volumes (µl) are shown in parentheses. From the Y/connector to the membrane start, the inlet tube is 75 mm (∼1.3 µl) and the outlet tube is 74.5 mm (∼1.2 µl). The connector block contributes 11.8 mm (∼0.1 µl). The membrane length is 3.0 mm (∼0.06 µl). Short shaft segments near the tip are indicated (5, 2.6, and 1.7 mm), with a small pre-membrane lumen of ∼0.2 µl. Inset: tip cross-section illustrating perfusate flow inlet to membrane region.

**Supplementary Figure S2:**
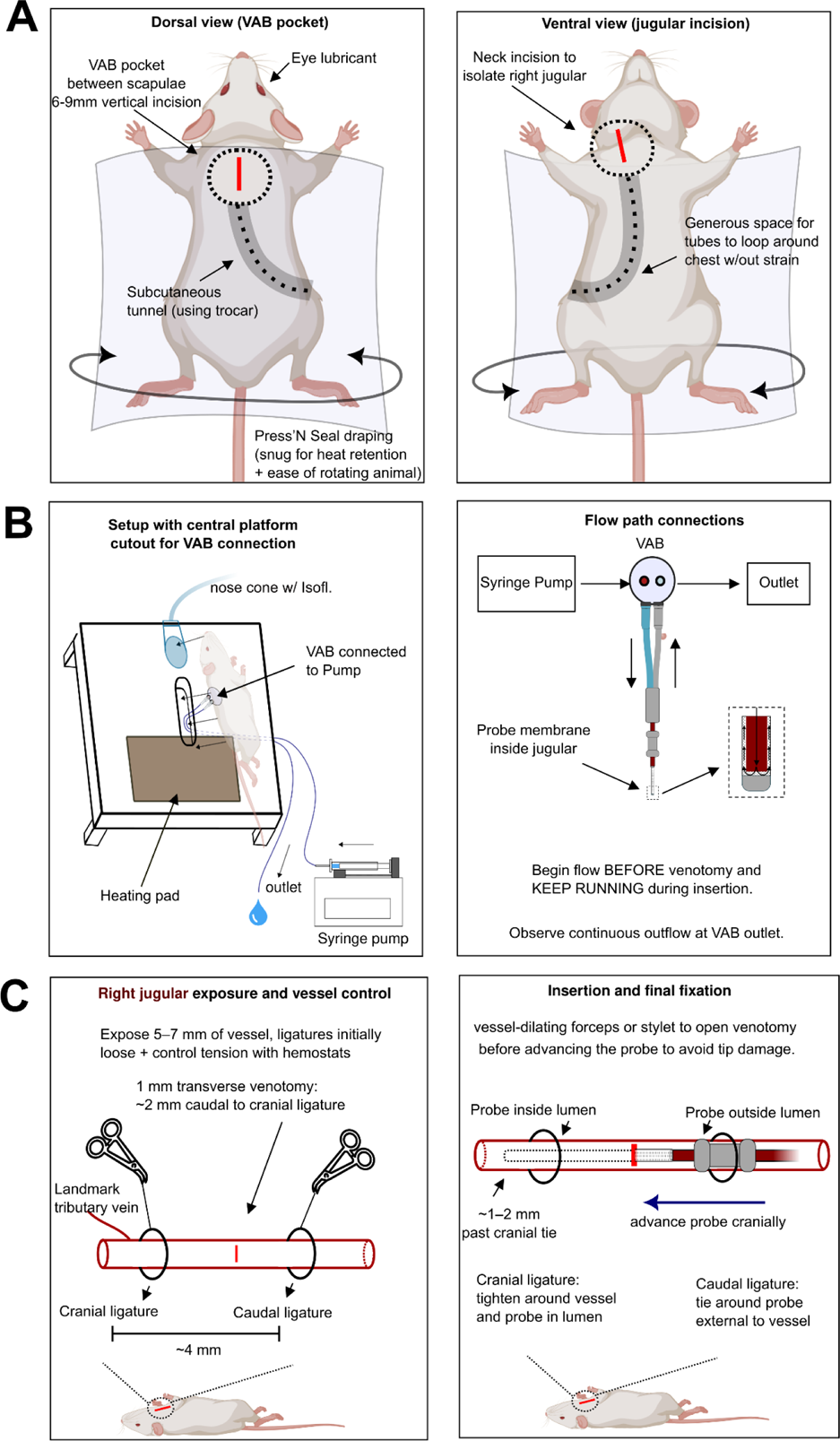
Vascular-access button–assisted jugular microdialysis in mouse. (A) **Left:** Mouse is draped with Press’N Seal (snug to retain heat and allow easy rotation). A 6–9 mm midline dorsal incision is made between the scapulae to form the VAB pocket. A subcutaneous tunnel is created using a trocar from the VAB pocket to the neck. **Right:** On the ventral side, a short neck incision is made to isolate the right jugular vein. Leave generous space for tubing to loop around the chest to provide strain relief. **(B) Left:** Setup on a heated platform with a central cutout that accommodates the VAB seated subcutaneously between the scapulae; nose cone delivers isoflurane. **Right:** Connect syringe pump → VAB inlet → probe → VAB outlet → collection, and begin flow before the venotomy; observe continuous outflow at the VAB outlet. **(C) Left:** Expose 5–7 mm of the right jugular. Identify the dorsolateral tributary as the cranial landmark. Place two 6-0 silk ligatures 3–4 mm apart, both initially loose, and apply opposing hemostats for gentle partial occlusion. Plan a 1 mm transverse venotomy ∼2–3 mm caudal to the cranial ligature. **Right:** Advance the probe cranially so the tip lies ∼1–2 mm past the cranial ligature inside the lumen. Tighten the cranial ligature around the vessel and probe (non-occlusive), then tie the caudal ligature around the probe external to the vessel. Keep flow running and verify continuous outflow.

**Supplementary Figure S3:**
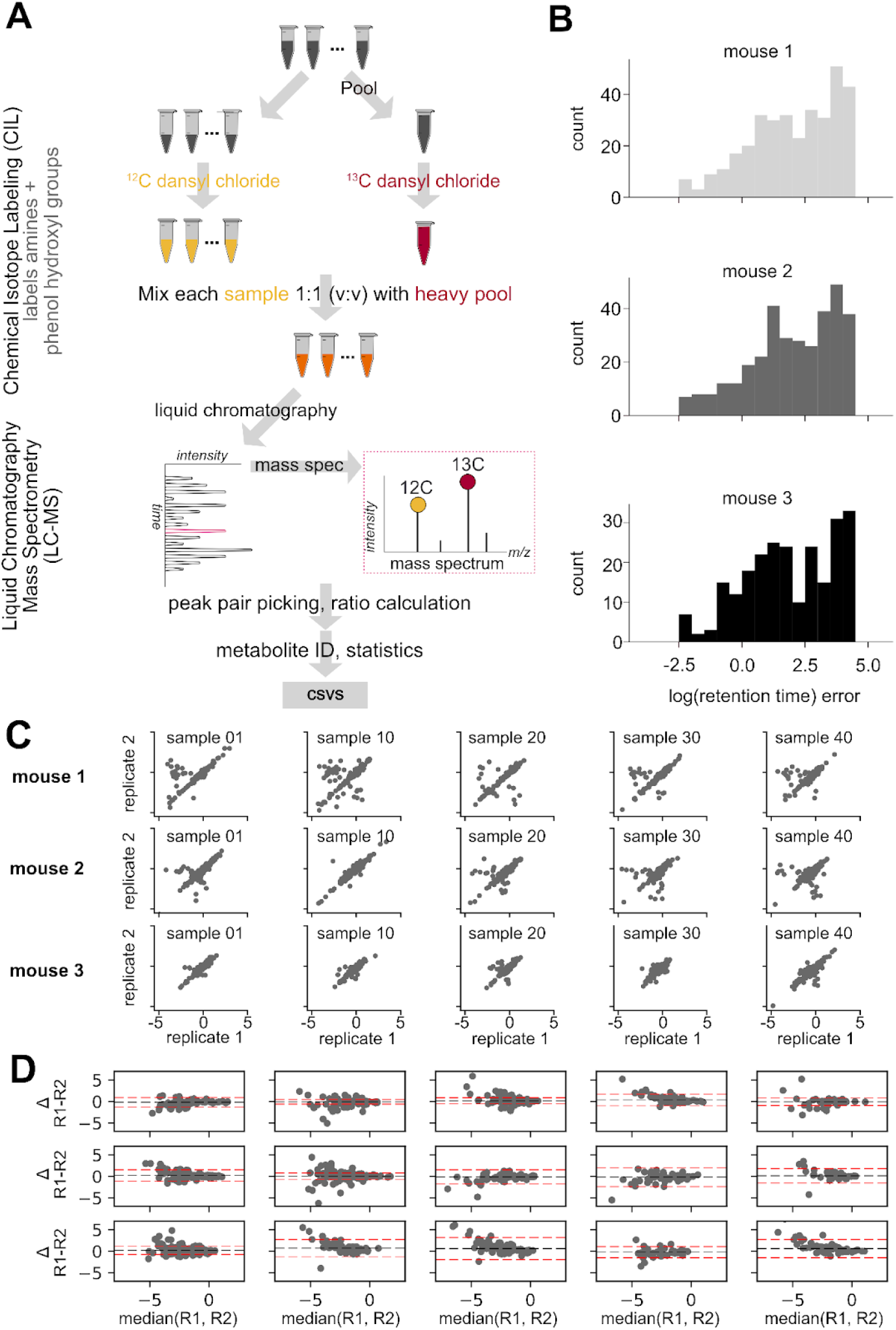
Chemical isotope labeling LC-MS workflow and analytical quality control. **(A)** Sample preparation and chemical Isotope labeling LC-MS workflow for amines and small peptides. **(B)** Drift-corrected retention-time (RT) error distributions per animal. **(C)** Technical replicate scatter plots (log scale). **(D)** Bland-Altman plots (difference vs. mean replicate); dashed lines represent 95% confidence interval (CI).

**Supplementary Figure S4:**
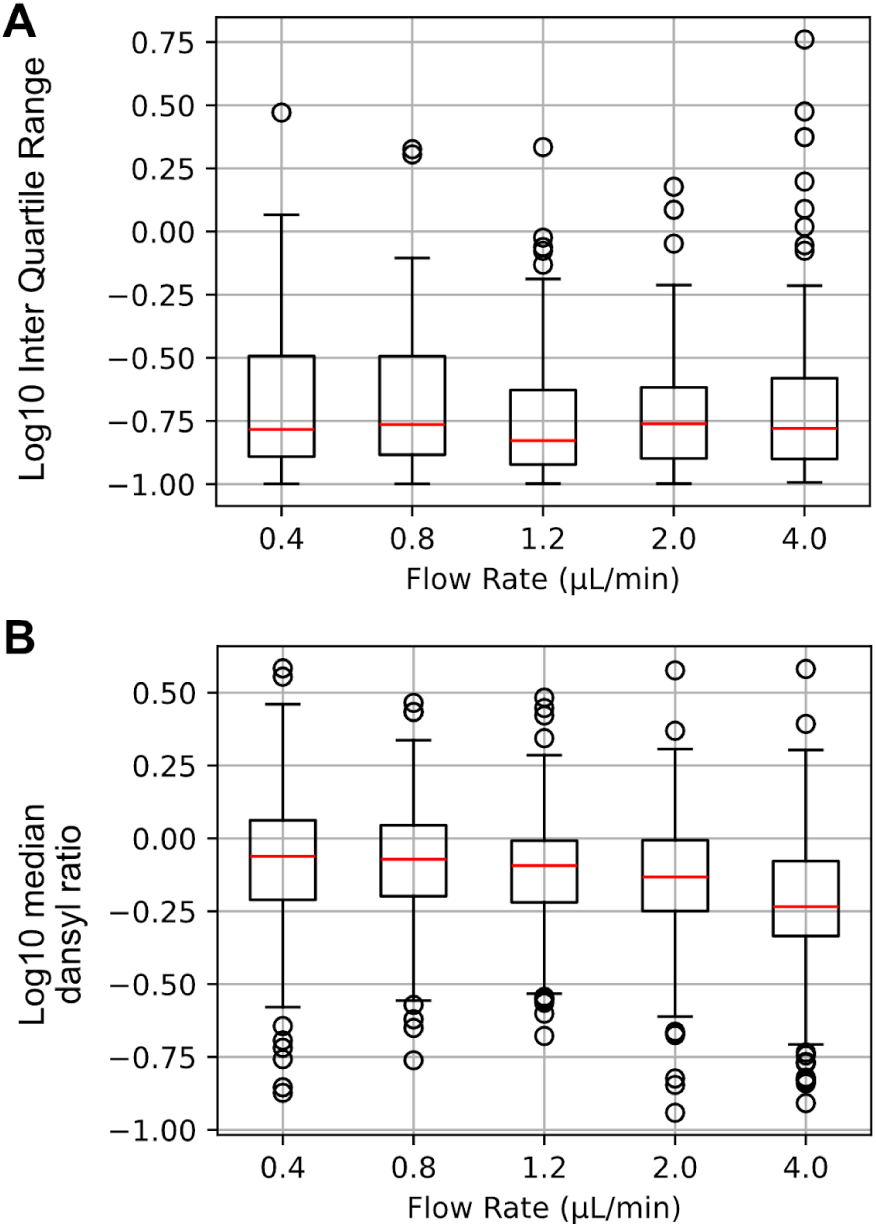
Relative recovery vs flow rate. Dialysate signal statistics across flow rates (0.4-4.0 µL/min) to analyze the tradeoff between recovery and temporal precision. For each flow rate, we collected 20µL samples, which were divided into three subsamples with two technical replicates each. **(A)** Boxplot for the distribution of interquantile range across 6 technical replicates. Lower is better. **(B)** Boxplot for the distribution of median dansyl ratio across the 6 replicates. Higher is better.

**Supplementary Figure S5:**
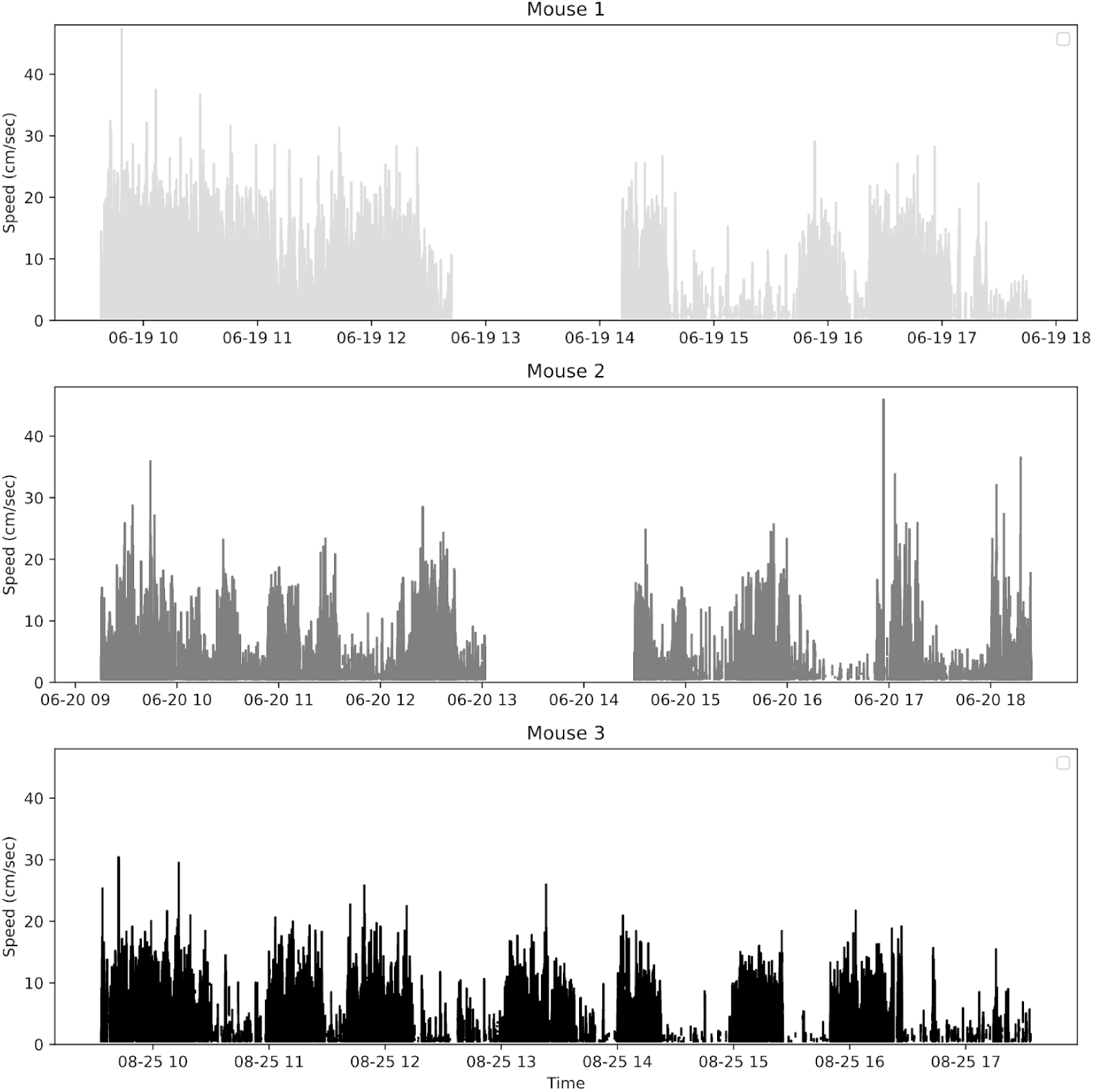
Mouse locomotion across hours of recordings in three mice during MD recordings. Speed traces over ∼8 h illustrate alternating bouts of movement and immobility that structure our physiology vs. behavior analyses.

**Supplementary Figure S6:**
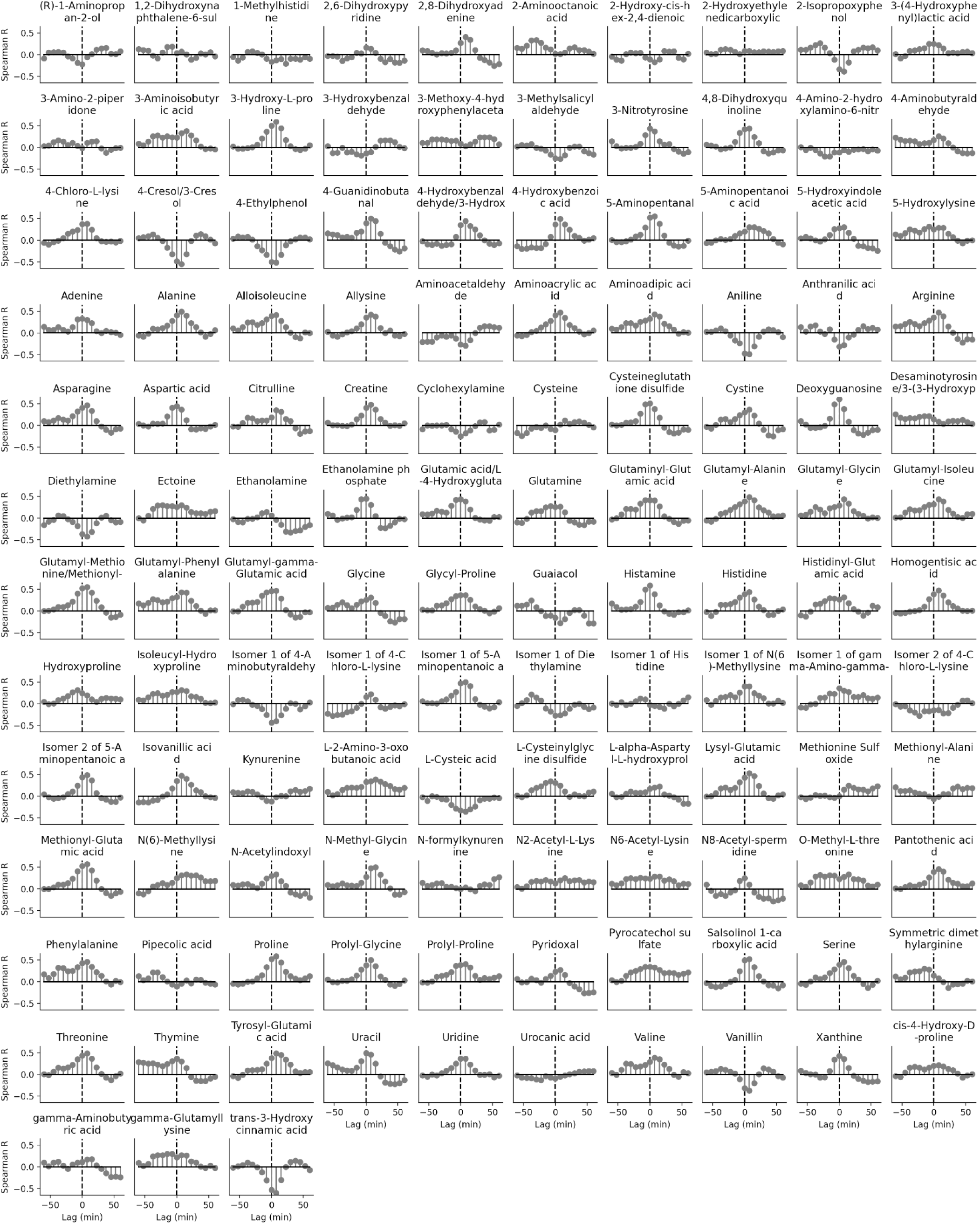
Grid of per-compound cross-correlations with locomotion across lags (−60 to +60 min, 7.5-min steps) for the 123-analyte high-quality intersection across animals.

**Supplementary Figure S7:**
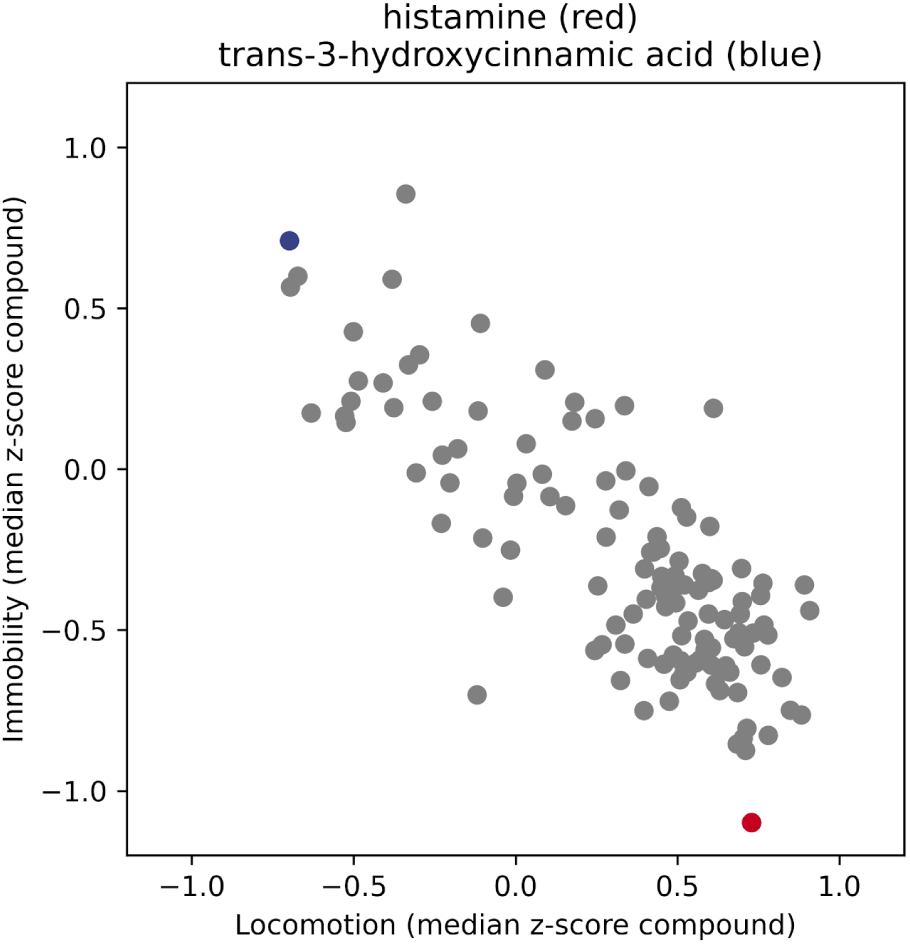
Immobility vs locomotion compounds. We selected two thresholds: med ± MAD/2 (med = median, MAD = median absolute deviation) of locomotion measured in 7.5 min bins, and define immobility where animal avg. locomotion was below med - MAD/2 for at least 2 time bins, and locomotion when the animal avg. locomotion was above med + MAD/2 for at least 2 time bins. Each point in the plot corresponds to the average z-scored value of each compound during periods of immobility vs locomotion, with red and blue highlighting Histamine and Hydroxycinnamic acid, respectively.

**Supplementary Figure S8:**
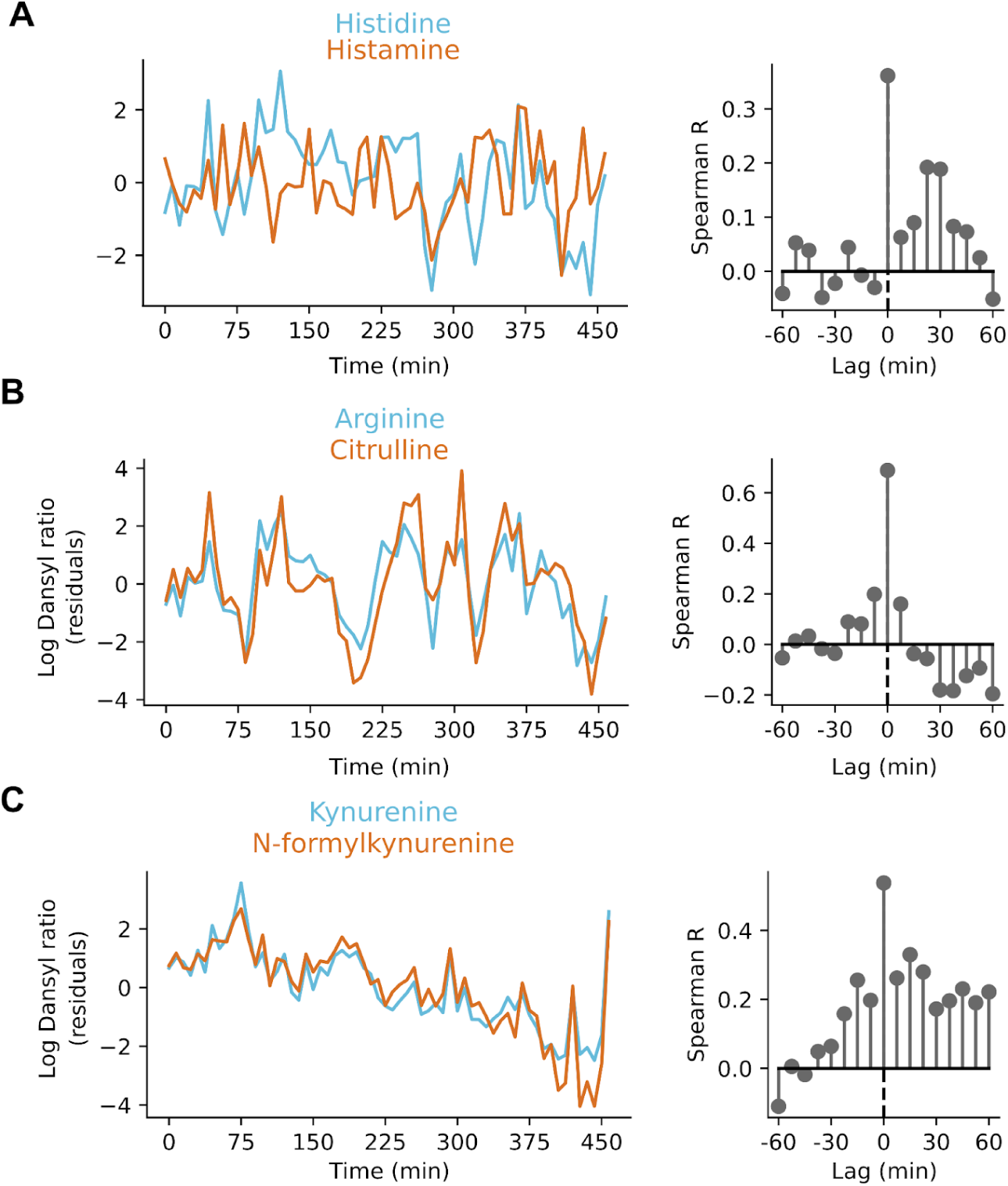
Interactions across compounds expected from literature. Analyses of locomotion-corrected compound timeseries interactions focusing on 3 literature-backed examples using data only from animal 3. **(A)** Histamine is produced from histidine through decarboxylation (71); **(B)** Nitric oxide synthase converts arginine to citrulline (72); **(C)** Arylformamidase converts N-formylkynurenine to kynurenine (73).

**Supplementary Figure S9:**
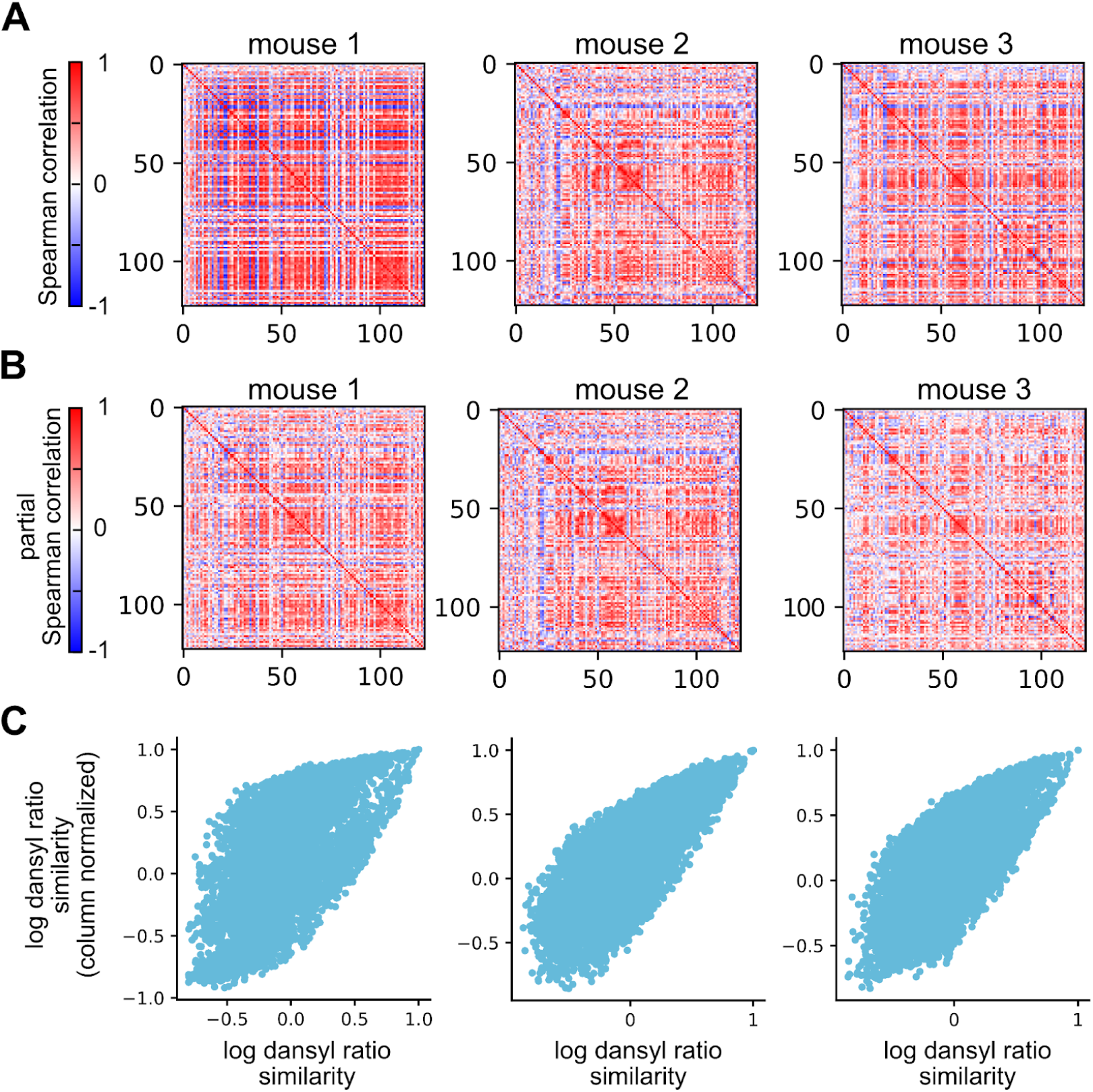
Stability of covariation structure across animals and controls. **(A)** Spearman correlation matrices per animal (Cosine similarity across matrices all > 0.5, p<1e-10). **(B)** Partial correlations controlling for locomotion (Cosine similarity across matrices all > 0.5, p<1e-10). **(C)** To rule out potential artifacts from time-varying recovery (e.g., due to blood flow or temperature), we z-scored the log-ratios of each sample. Sample-wise normalization preserves the correlation structure (each animal’s Spearman’s rank correlation coefficient > 0.8, p<1e-10). Scatterplot for the three animals’ original vs z-scored compound similarity is shown.

**Supplementary Figure S10:**
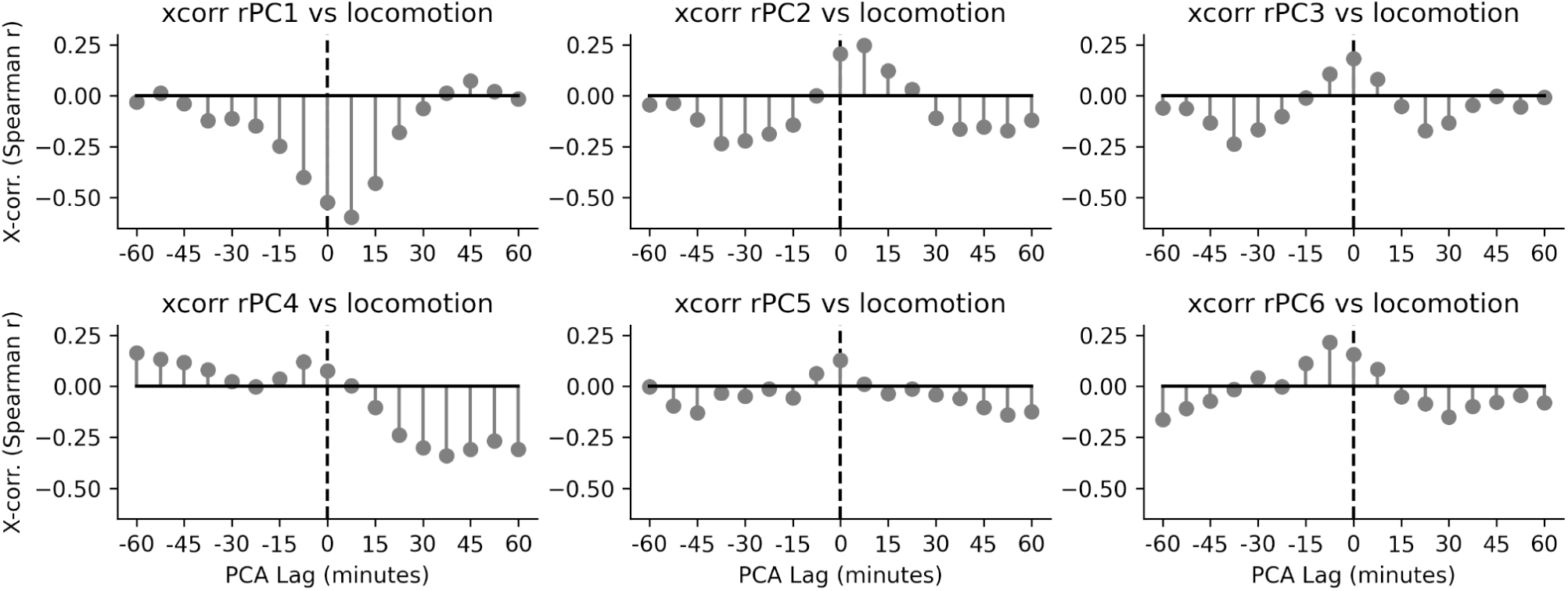
Behavior alignment across principal components. Cross-correlations of PCs with locomotion (−60 to +60 min) show the strongest positive alignment for rPC1 with a 7.5 min positive lag.

## Supplementary Tables

**Supp. Table 1:** Comparison of methods for blood profiling during behavior. Attributes of invasive sampling, non-invasive byproducts, biochemical/electrochemical sensors, and microdialysis. Across rows, we focus on the medium sampled; the time resolution of the method used, such as the delay of the measurement compared to the actual circulating compound; the frequency, as in how often the sampling can be made; the number of compounds that can be measured/recovered with that technique; and some key references. Microdialysis offers minute-scale cadence, high multiplexing, and compatibility with freely moving animals.

|  | <u>Blood sampling -- vein pricks</u> | <u>Blood sampling -- vascular catheter</u> | <u>Byproducts and external analyses</u> | <u>Biochemical / electrochemical sensors</u> | <u>Microdialysis</u> |
| --- | --- | --- | --- | --- | --- |
| Medium | *blood* | *blood* | *urine, feces, saliva, sweat, genital swab* | *CSF, blood* | *CSF, blood, tissue, plasma* |
| Time Resolution | sec - min | sec - min | min-hrs | ms - s | min |
| Frequency | hrs<sup>-1</sup>/day<sup>-1</sup> | hrs<sup>-1</sup> | min<sup>-1</sup>/h<sup>-1</sup> | s<sup>-1</sup> | min<sup>-1</sup> |
| Number of Compounds | 1000s | 1000s | 100s | 10s | 100s |
| Key References | (17) | (19) | (15, 83–85, 95) | (15, 96, 97) | (21, 22, 26, 30, 32, 44, 98) |

**Supp. Table 2:**
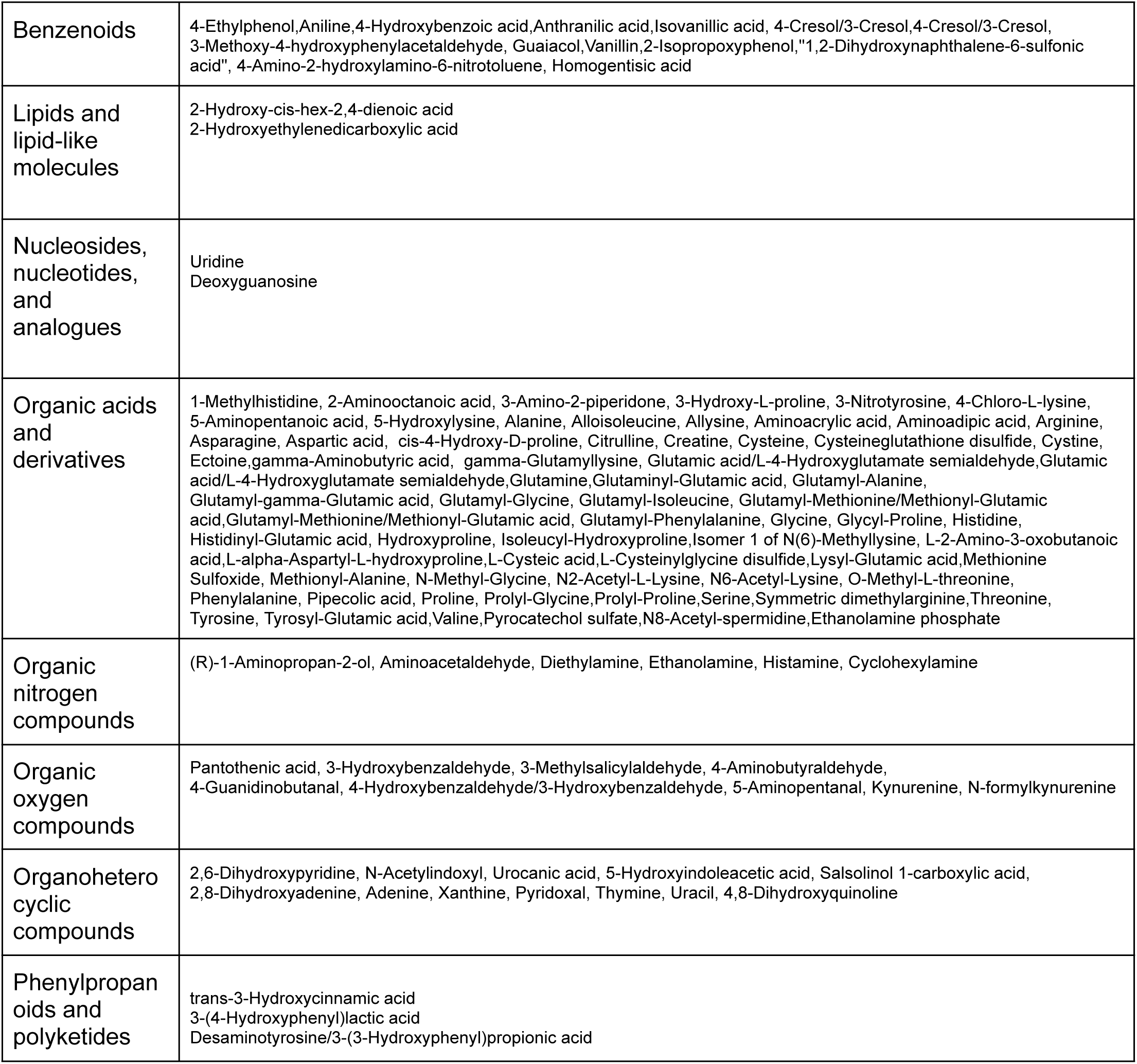
High-quality intersection compounds used for manifold analyses. ChemOnt taxonomy class (99) determined according to PubchemID and automated chemical classification *ClassyFire* (99) available at https://cfb.fiehnlab.ucdavis.edu/.

